# Extracellular Vesicles Analysis in the COVID-19 Era: Insights on Serum Inactivation Protocols Towards Downstream Isolation and Analysis

**DOI:** 10.1101/2020.12.10.417758

**Authors:** Roberto Frigerio, Angelo Musicò, Marco Brucale, Andrea Ridolfi, Silvia Galbiati, Riccardo Vago, Greta Bergamaschi, Anna Ferretti, Marcella Chiari, Francesco Valle, Alessandro Gori, Marina Cretich

**Affiliations:** Istituto di Scienze e Tecnologie Chimiche “Giulio Natta” (SCITEC) - Consiglio Nazionale delle Ricerche; Istituto per lo Studio dei Materiali Nanostrutturati (ISMN) - Consiglio Nazionale delle Ricerche; Consorzio Interuniversitario per lo Sviluppo dei Sistemi a Grande Interfase (CSGI), Florence, Italy; IRCCS San Raffaele Scientific Institute, Milano, Italy

## Abstract

Since the outbreak of COVID-19 crisis, the handling of biological samples from confirmed or suspected SARS-CoV-2 positive individuals demanded the use of inactivation protocols to ensure laboratory operators safety. While not standardized, these practices can be roughly divided in two categories, namely heat inactivation and solvent-detergent treatments. As such, these routine procedures should also apply to samples intended for Extracellular Vesicles (EVs) analysis. Assessing the impact of virus inactivating pre-treatments is therefore of pivotal importance, given the well-known variability introduced by different pre-analytical steps on downstream EVs isolation and analysis. Arguably, shared guidelines on inactivation protocols tailored to best address EVs-specific requirements will be needed among the EVs community, yet deep investigations in this direction haven’t been reported so far.

In the attempt of sparking interest on this highly relevant topic, we here provide preliminary insights on SARS-CoV-2 inactivation practices to be adopted prior serum EVs analysis by comparing solvent/detergent treatment vs. heat inactivation. Our analysis entailed the evaluation of EVs recovery and purity along with biochemical, biophysical and biomolecular profiling by means of Nanoparticle Tracking Analysis, Western Blotting, Atomic Force Microscopy, miRNA content (digital droplet PCR) and tetraspanin assessment by microarrays. Our data suggest an increase in ultracentrifugation (UC) recovery following heat-treatment, however accompanied by a marked enrichment in EVs-associated contaminants. On the contrary, solvent/detergent treatment is promising for small EVs (< 150 nm range), yet a depletion of larger vesicular entities was detected. This work represents a first step towards the identification of optimal serum inactivation protocols targeted to EVs analysis.

## Introduction

The COVID-19 pandemic forced researchers to deal with clinical specimens from confirmed or suspected SARS-CoV-2 positive cases. Current biocontainment guidelines to address lab operators exposure risk are adopted according to international standards and constantly updated (https://www.cdc.gov/coronavirus/2019-nCoV/lab/lab-biosafety-guidelines.html). In this regard, the minimum biosafety level to handle suspect SARS-CoV-2 specimens during non-propagative procedures is BSL-2, provided that the samples have been biologically inactivated to abolish or mostly suppress virus infectivity. Common inactivation protocols are inherited from previous studies on enveloped viruses validated during the past MERS or SARS outbreaks; those preceding molecular diagnostics (involving RNA extraction) are usually based on chemical treatments with detergents and chaotropic agents^1, 2^. Previous experience on serology of coronaviruses also suggested treatments with a solvent-detergent combination (e.g. Triton X100/Tween 80 and tri(n-butyl) phosphate), as currently adopted for serum/plasma standards by the Medicine & Healthcare products Regulatory Agency^3^. Heat treatment is another routine inactivation method, especially for serum/plasma. On this matter, while data are still debated^1, 4, 5^ serum heat inactivation at 56°C for 30 min is emerging as a common practice.

In this scenario, arguably, it is anticipated that assessing the impact of different serum inactivation protocols on downstream Extracellular Vesicles (EVs) isolation and analysis will be of primary relevance to the EVs community while, to the best of our knowledge, no investigation has been reported so far in this direction. EVs from biological samples are indeed tremendously complex analytes, and the well-known influence of pre-analytical practices on downstream EVs use has been driving the need for standardization and rigor criteria largely before the COVID crisis, as highlighted by the ISEV community^6, 7^.

Herein, we report on the influence of two COVID-19 serum inactivation protocols on EVs recovery, purity, biophysical, biochemical and biomolecular traits. Specifically, on unbiased premises, we focused our investigation on EVs isolated by ultracentrifugation (UC) from untreated (NT), heat treated (HT) and solvent/detergent (S/D) treated healthy sera. Samples were analyzed by means of Nanoparticle Tracking Analysis (NTA), Western Blotting (WB), Atomic Force Microscopy (AFM), miRNA 16-5p and miRNA 21-5p quantification by droplet digital PCR (ddPCR) and antibody microarrays for tetraspanin assessment. A flow chart of the experimental strategy is reported in Scheme 1. Our data showed that serum heat inactivation provided the highest UC recovery, yet inclusive of contaminants enrichment. Solvent/detergent treatment leads to no remarkable effect when small EVs (< 150nm range) are considered, providing the best EVs purity among the three groups. Far from being conclusive, our work aims to provide preliminary insights and awareness among the EVs-community on viruses inactivation practices to be adopted prior serum EVs analysis,

**Scheme 1.**
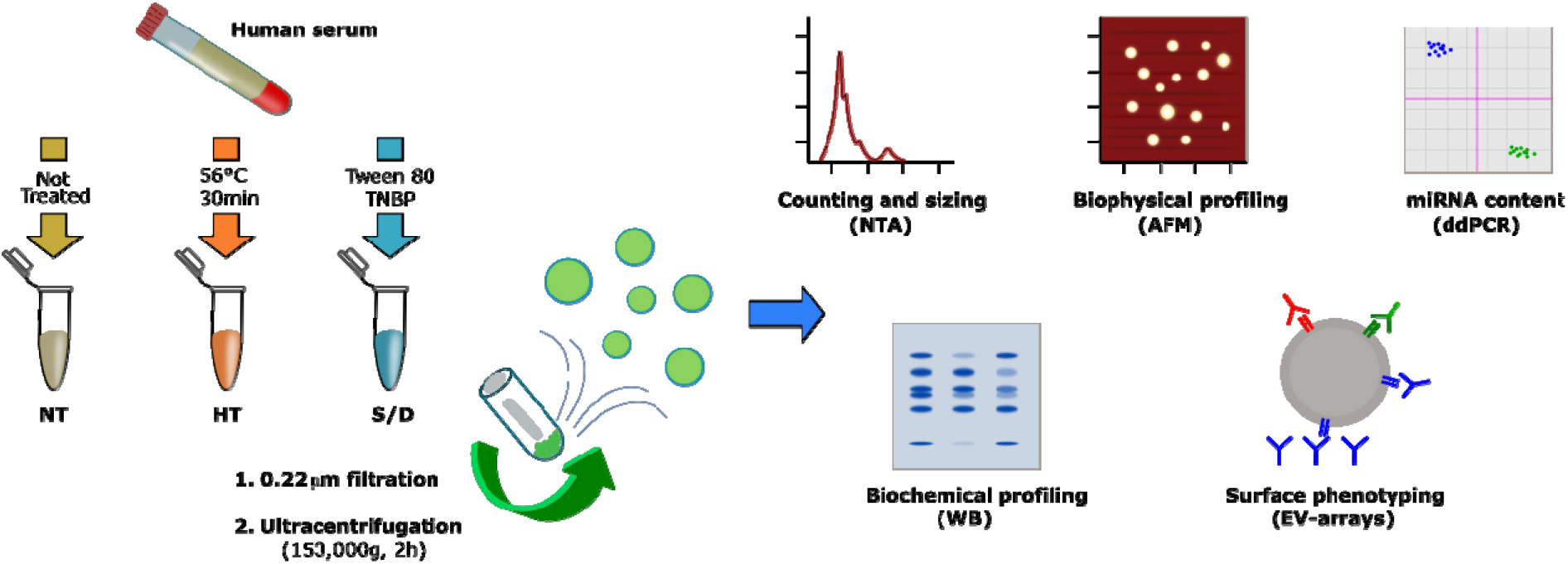
Workflow describing the sample treatments, EV isolation and characterization

## Results and discussion

### Sample preparation

Sixteen pre-COVID serum samples from healthy donors were divided in three aliquots (750uL each). For each serum sample, one aliquot was left untreated (NT), one aliquot (HT) was set to mimic heat inactivation (56°C for 30 minutes), and one aliquot (S/D) was treated with 10mg/mL Tween 80 and 3 mg/mL tri(n-butyl) phosphate (TNBP). Hints on compatibility of such treatments with EV integrity were previously reported in studies on the stability of vesicles upon different temperatures^8^ and non ionic detergents^9^. The resulting 48 samples were subjected to standard ultracentrifugation (UC) at 150.000g for 2 hours. It is well documented that single-step EVs isolation procedures, including UC, are likely to lead to EVs co-isolation of contaminants such as protein aggregates, VLDLs, LDLs and chylomicrons^10, 11^, whereas a combination of sequential purification steps provide increased purity^12 13^. As such, we reasoned that the simple and routinely performed EVs isolation by UC could be particularly indicative in assessing the role of serum pretreatment on the extent of co-isolated contaminants.

### Nanoparticle Tracking Analysis

Pellets from UC samples were resuspended in PBS (50μL), and particle number and sizing of the 48 samples were determined by Nanoparticle Tracking Analysis (NTA) as described in Methods section. The resulting particle concentration (A), mean (B) and median (C) of particle diameter for the untreated (NT), heat treated (HT) and solvent/detergent treated (S/D) samples are shown in Figure 1.

**Figure 1:**
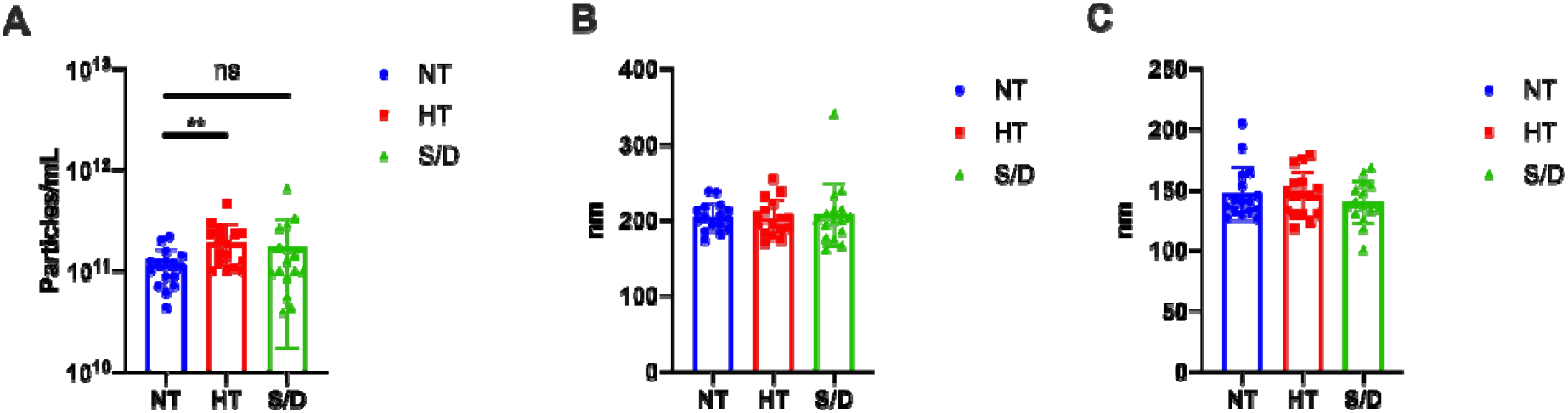
NTA analysis of EVs isolated by ultracentrifugation from untreated (NT), heat treated (HT) and solvent/detergent (S/D) treated healthy sera. N= 16. A: mean particle count. B: mean particle size. C: median particle size. Significative: p<0.05; * = p<0.05; ** = p<0.01.

NTA analysis revealed a significant (p < 0.01) increase in particle counting in the HT EVs compared to the NT sample, whereas no difference was detected at a statistically significant level for the S/D treated EVs (Figure 1A). Yet, the S/D samples show a higher variability in the number of UC recovered particles (Figure 1A). As for particle mean and median size, no significant differences were found among the three sample sets. It is well known that given the presence of co-isolated lipoproteins, the quantification of EVs based on particle counting by NTA tends to overestimate EVs concentration^14^. Thus, we preliminary hypothesized that the increased number of recovered particles that is observed after heat-treatment could be ascribed to the increased co-precipitation of lipoproteins and other proteins aggregates triggered by heat-induced aggregation.

### Western Blotting

Western blotting (WB) was used to confirm the presence of EVs transmembrane (CD9 and CD63) and luminal proteins (Alix and TSG101), as well as the presence of common co-isolated contaminants (Apolipoprotein A I, Apo AI) in a set of NT, HT and S/D samples. Prior to WB, protein concentration in the UC-recovered pellets was assessed by Bradford assay showing a remarkably higher protein content in the HT sample (10.6 mg/mL vs. 3.8 mg/mL for both NT and S/D). The samples were then diluted and loaded on the gel at the same protein amount per lane (5 ug).

Figure 2 shows the results of the WB gels for 2 representative samples for each sample group, analyzed in non-reducing conditions (A), reducing conditions (B) and the corresponding immunoblotting for the assessment of TSG101 (C), Alix (D), CD9 (F), CD 63 (G). Overall, the presence of typical EVs markers was demonstrated for all the three sample sets with similar isolation yields. However, a higher amount of coisolated Apo AI is clearly detectable in the HT group (E).

**Figure 2.**
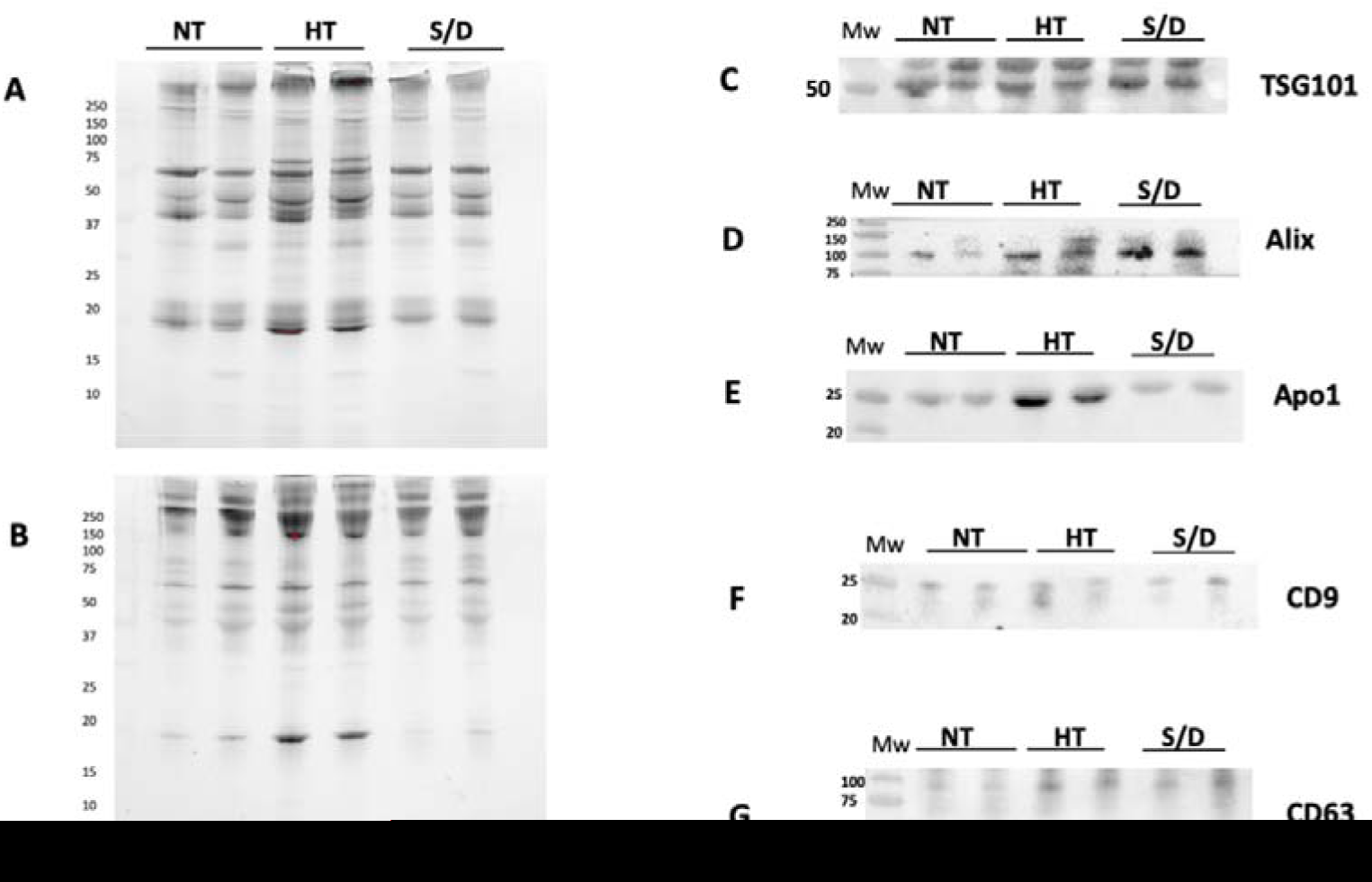
Western Blotting analysis of EVs isolated by ultracentrifugation (UC) from untreated (NT), heat treated (HT) and solvent/detergent (S/D) treated healthy sera. N= 12. The SDS PAGE of pellets was run in in non-reducing conditions (A) and reducing conditions (B). Immunoblotting was performed for TSG101 (C); Alix (D); contaminant Apoliprotein AI (E); tetraspanin CD9 (F) and CD63 (G).

The quantification of intensity for each immune-blotted protein band was performed and averaged. We then calculated the ratio between EV-specific protein markers and the co-isolated Apo AI contaminant in order to estimate and compare the purity yield of isolated EVs after each inactivation treatment. Results are reported in Figure 3. An increase of co-isolated lipoproteins following heat treatment is clearly observable (p < 0.01) by comparing the ratio between the luminal marker TSG101 and Apo AI in the three groups. The same trend was observed considering the ratio between ALIX and Apo AI. On the other hand, no clear difference among the samples is detectable when the ratio between tetraspanins CD9/CD63 and Apo AI is considered. The selection of appropriate markers for such comparison, as a consequence, may prove extremely critical, posing multimarker selection as likely mandatory. Overall, an apparent reduction in lipoprotein contaminants was observed in the case of S/D treatment, even in comparison with untreated samples. This observation suggests a role of solvent/detergent in shielding those supramolecular interactions at the colloidal level among EVs and lipoproteins, that may contribute to co-isolation.

**Figure 3.**
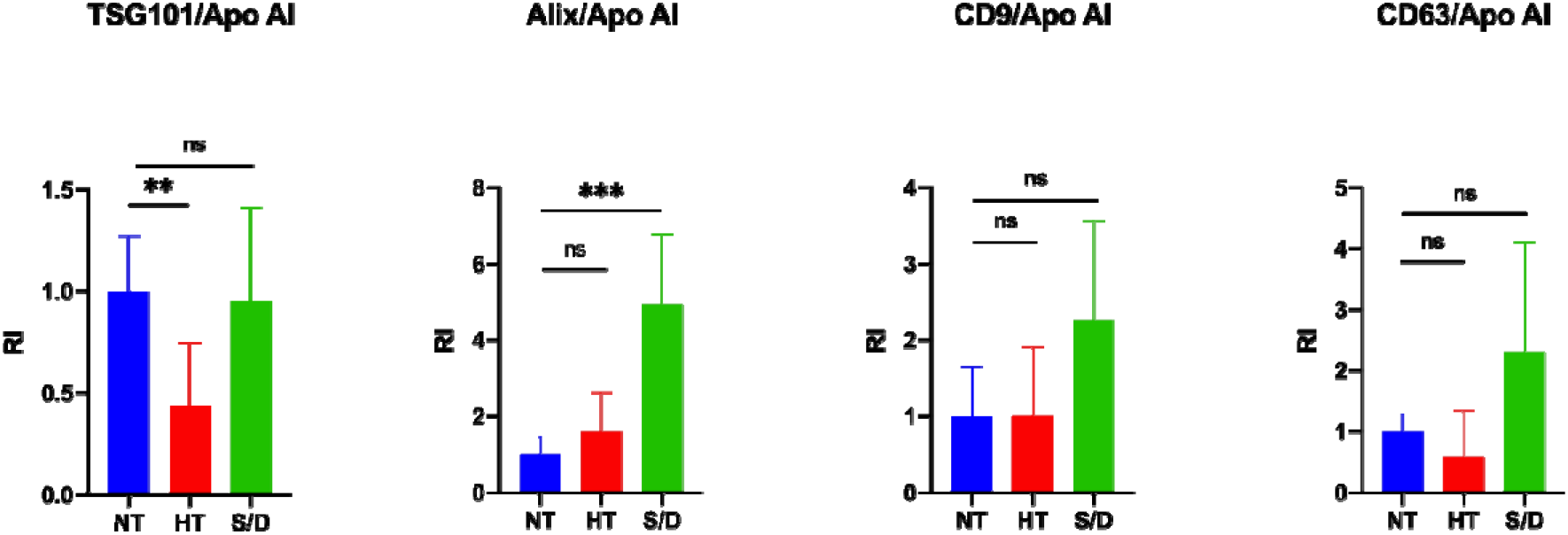
Quantification of blotted protein bands and ratio between EV luminal markers TSG101/Alix and EV surface markers CD9CD63 with contaminant lipoprotein Apo AI. Significative: p<0.05; * = p<0.05; ** = p<0.01; *** = p<0.001.

### Atomic force microscopy (AFM)

Samples collected from different serum inactivation protocols were analyzed via a high-throughput nanomechanical screening method described elsewhere^15^. Briefly, the vesicle/surface contact angle (CA) of individual EVs adsorbed on a substrate can be measured via AFM morphometry and used as a direct indication of their mechanical stiffness. The same procedure allows calculating the diameter of each observed EV prior to surface adsorption. Vesicular objects are characterized by a narrow distribution of CAs at all diameters, whereas non-vesicular contaminants show a wider dispersion of CAs^16^ which can be used to infer their presence in a sample even when their globular morphology makes it difficult to discern them from EVs. Figure 4 summarizes the main differences revealed by AFM morphometric analysis across the panel of samples.

**Figure 4.**
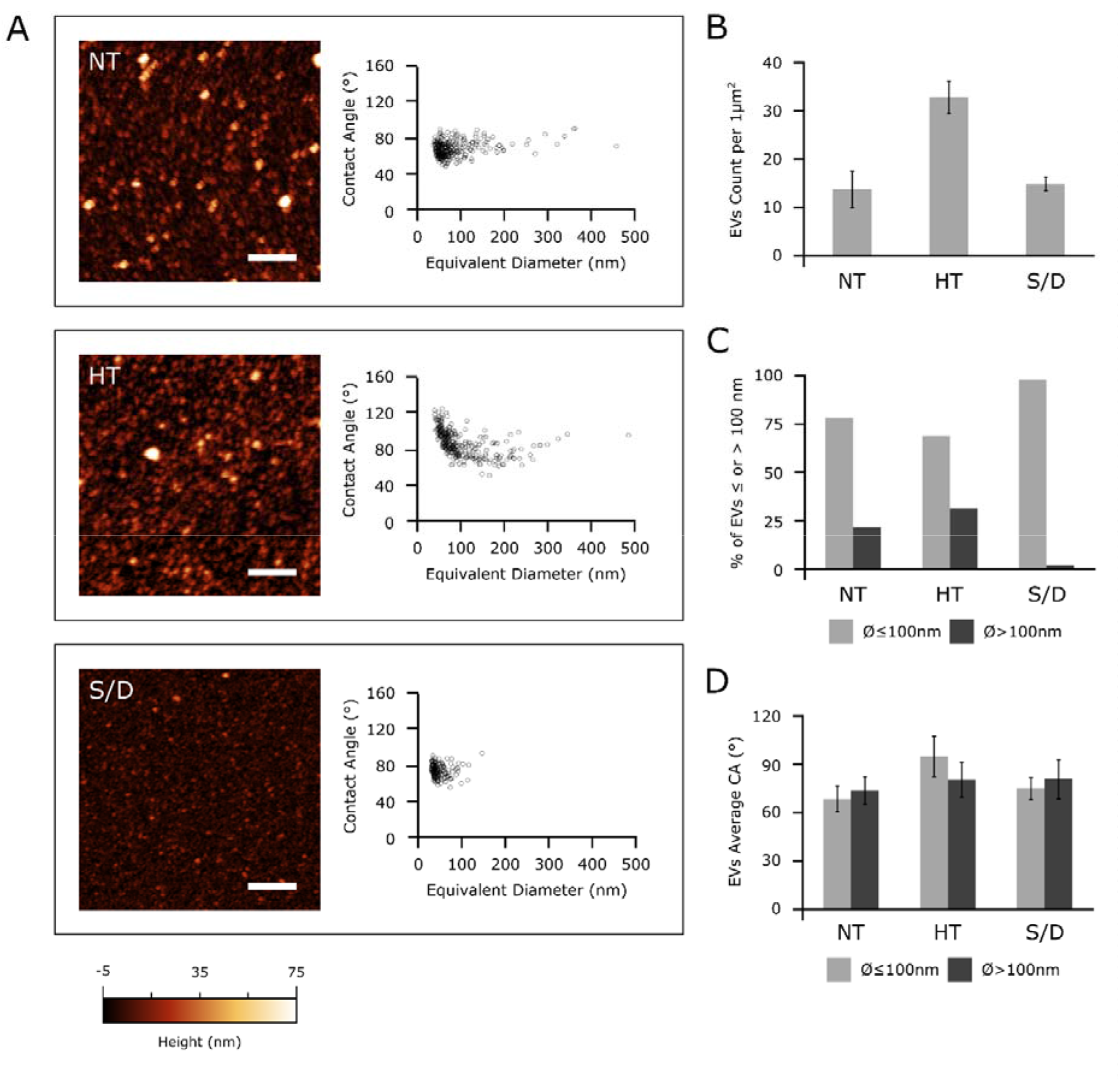
(A): left column – representative AFM micrographs of NT, HT and S/D samples. Scale bars are 1μm. Right column – scatterplots of surface contact angle VS diameter in solution of EVs measured via quantitative AFM morphometry as described elsewhere.^15^ Each circle represents an individual EV. (B): surface density of globular objects in NT, HT and S/D samples deposited with the same protocol (see materials and methods); (C) percentage of adsorbed EVs having diameters above or below 100 nm in their spherical conformation; (D) average surface/vesicle contact angle (representative of mechanical stiffness) of EVs with diameters above or below 100 nm.

All examined EVs samples showed an abundant vesicular content (Figure 4A, left column); however, in accordance with the increased particle counting observed via NTA (Figure 1), the HT sample showed more than twice the amount of adsorbed globular objects (Figure 4B) with respect to NT and S/D samples deposited with the same procedure (see materials and methods).

Figure 4A shows the CA vs diameter plots of around 200-300 individual EVs for each sample. All three samples were found to contain a high proportion of globular objects with diameters in the 50-100 nm range and a decidedly smaller amount of objects with diameters between 100 and 500 nm. In particular, only 2% of the EVs in the S/D sample had a diameter above 100nm (Figure 4C), with no individual S/D treated EV having a diameter >150 nm (Figure 4A). In contrast, a more substantial amount of EVs in both the other samples had diameters above 100nm (respectively 22% and 31% of the EVs measured in NT and HT samples, Figure 4C).

Figure 4D shows the average CAs of EVs smaller and larger than 100 nm in the three samples. The NT CA/diameter scatterplot in Figure 4A does not show any significant CA discontinuity between the two ranges of diameters; accordingly, average CA values of smaller (68±8°, N=242) and larger (73±8°, N=67) EVs are similar in this sample (Figure 4D). The S/D sample shows very similar values (74±7°, N=171 for smaller and 79±12°, N=4 for larger EVs), suggesting that while the solvent/detergent treatment dramatically reduced the amount of larger EVs, it did not significantly impact the structural integrity of the remaining EVs, which continue to show the same mechanical characteristics of untreated ones. The same consideration can be made for larger EVs in HT samples (average CA = 79±11°, N=103). In contrast, globular objects with diameters below 100nm have a significantly higher CA (94±12°, N=226) in HT, suggesting marked structural or compositional differences in this sub-population of objects with respect to other samples.^12, 17^ Taken together, these results suggest the possibility that NT samples might contain different types of vesicular objects sharing similar mechanical characteristics but having different average dimensions, with a diameter threshold of around 100 nm separating the main subpopulations.

Although the absence of significant CA differences across all sizes in non-treated EVs makes this hypothesis only tentative, HT and S/D treatments seem to selectively act on only some of the putative subpopulations: S/D is able to deplete larger EVs, while HT enriched the solution with a population of objects with distinct mechanical properties.

### Microarray analysis

EVs microarrays are high-throughput analytical platforms that are used to phenotype EVs. In this technique, antibodies^18^ or peptide ligands^19^ are used to capture EVs by their most common surface-associated proteins or by membrane sensing, followed by fluorescence immune-staining of membrane biomarkers. Here, the BP membrane binding peptide^19^ – that captures EVs by a general, membrane-mediated mechanism not involving surface antigens – was spotted on a silicon based microarray platform for enhanced fluorescence detection^20, 21^. EVs from the NT, HT and S/D groups were then incubated at 10^9^ particles/mL, and a cocktail of biotinylated anti CD9/CD63/CD81 followed by Cy3 labelled streptavidin was used for detection. Figure 5 reports the resulting fluorescence intensities for each group of samples. Results show a decrease of immune-reactivity particularly pronounced in the HT inactivated samples, that could suggest a lower content in tetraspanin-responsive particles; on the other hand we cannot rule out a possible partial denaturation of EVs surfaces markers affecting immune-staining. Overall, samples treatment appears to not preclude the possibility of EVs immune-phenotyping but the overall picture could prove tricky to be unambiguously defined.

**Figure 5.**
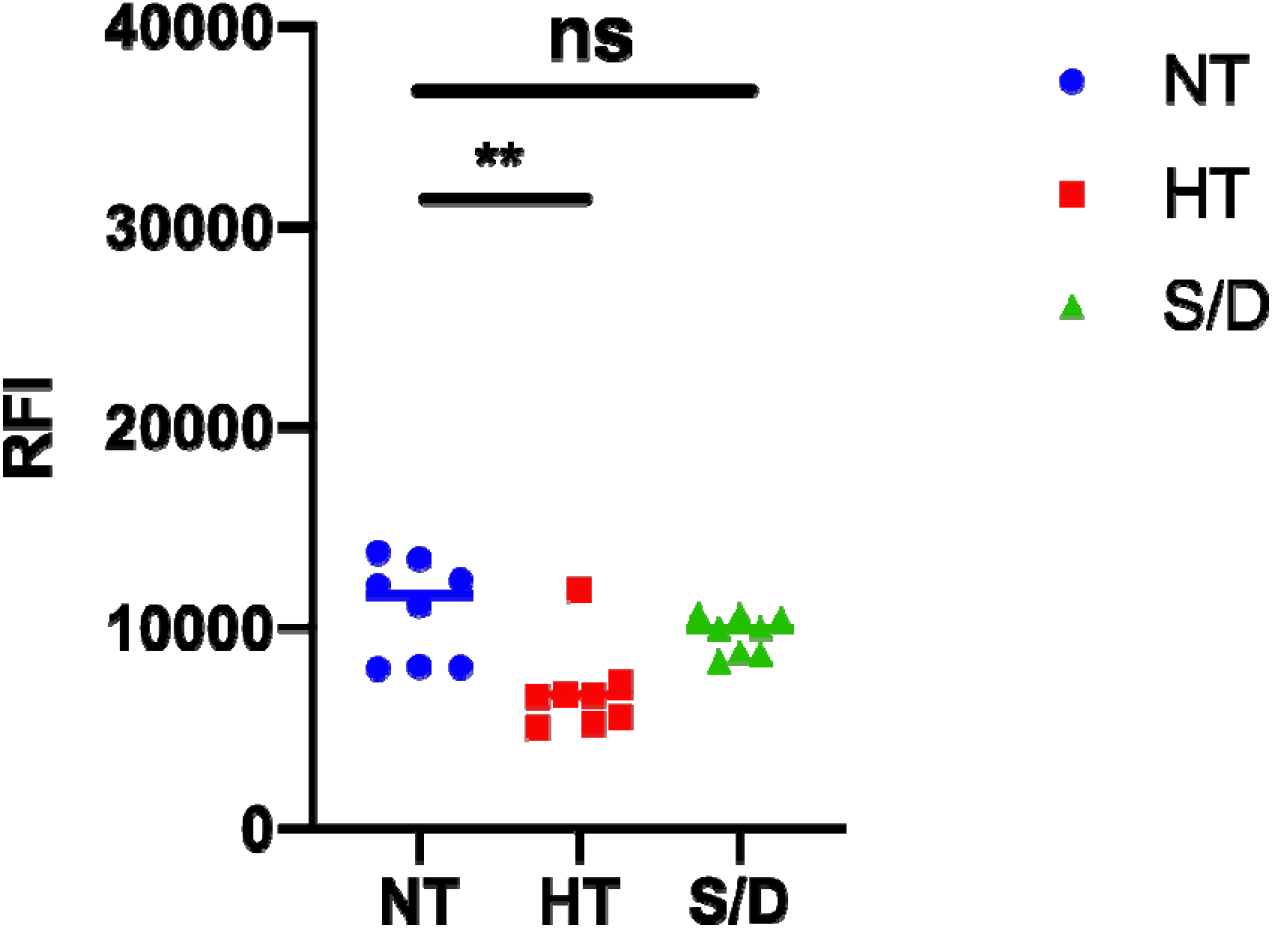
Results of immune-phenotyping by peptide microarrays of EVs isolated by ultracentrifugation from untreated (NT), heat treated (HT) and solvent/detergent (S/D) treated sera. EVs were captured by BP membrane binding peptide and fluorescently stained by a mixture of anti-CD9/CD63/CD81 antibodies. Significative: p<0.05; * = p<0.05.; ** = p<0.01

### miR-16-5p and miR-21-5p ddPCR analysis

Among different RNA classes, microRNAs (miRNA) are abundantly harbored in many body fluids via encapsulation and/or association to EVs as well as transported by lipoproteins^22^, which avoid nucleolytic degradation,.

We selected two representative miRNAs, namely miR-16-5p and miR-21-5p to compare if /how serum inactivation protocols could influence the miRNA levels.

MiR-16-5p is among the most abundant miRNA in EVs^23^ while miR-21-5p is reported to be associated to lipoproteins and could be involved in lipid metabolism^24^. We quantitatively detected miRNAs levels within NT, HT and S/D groups by ddPCR analysis; results are summarized in Figure 6. Data analysis highlights a clear trend for both miR-16-5p and miR-21-5p, with an increased amount detected within the HT-samples. This observation is consistent with the apparent higher EVs isolation yield for the HT samples suggested by protein quantification and NTA. On the other hand, given the reported data on EVs purity (see WB section) that instead suggested HT samples to contain higher levels of co-isolated contaminants, a more comprehensive interpretation should take into account the reported association of RNAs also to RNA-binding proteins (RBPs) and, especially for miRNA 21-5p, to high- and low-density lipoproteins^22, 25^.

**Figure 6.**
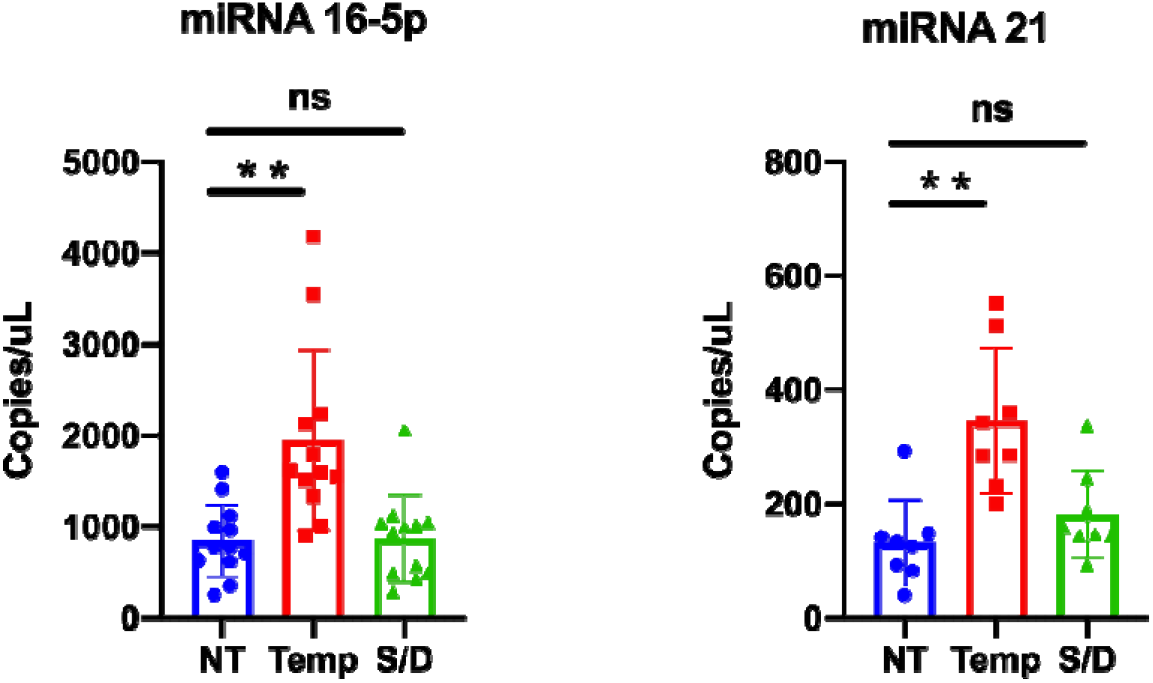
miR-16-5p and miR-21-5p expression levels in untreated (NT), heat treated (HT) and solvent/detergent (S/D) treated healthy sera analyzed by droplet digital PCR. Significative: p<0.05; * = p<0.05; ** = p<0.01.

## Conclusions

Our study was aimed at introducing preliminary insights on the role of different SARS-CoV2 inactivation protocols prior to serum EVs isolation and analysis. Exacerbated by the current pandemic scenario, this topic, arguably, will gain broad relevance to the EVs community. Clinical samples preparation is indeed known to have a profound impact on the isolation of EVs and related contaminants^6^. A full awareness on the effect of any additional sample pre-analytical treatment should be consequently arisen among EVs-users, particularly due to the fact that an interlaboratory consensus on protocols for SARS-CoV2 serum inactivation protocols is far from being reached. In this sense, laboratories that use bio-banked samples collected by clinicians may be particularly affected.

Far from being conclusive, our data suggest that the use of solvent/detergent addition could be seen a as a preferable virus deactivating method, taking EV’s purity obtained after a single UC step into account and as far as small EVs isolation and analysis are concerned. Non-ionic detergents are indeed relatively mild and usually non-denaturing^14^, yet known to break lipid-lipid and lipid-protein interactions. This could account for the apparent higher purity of EVs obtained after S/D treatment (see WB analysis), which contrasts with the higher content of lipoparticle contaminants detected following heat treatment. In this sense, their use upfront UC cycles could be worth of further investigation. On the other side, solvent/detergent treatment led to the depletion of vesicular particles of larger size (>150nm), and their use should be cautiously pondered if this EVs subpopulation represents the target of analysis. In contrast, heat-based protocols provided higher recovery, and the enrichment in EVs contaminants could be counterbalanced by the introduction of a subsequent step of purification. Overall, the virus inactivation procedure should be tailored considering the downstream analysis to be undertaken, and further work will be needed in this direction to identify the best possible practices.

## Materials and methods

### Ultracentrifugation

750 ul of serum were diluted 1:1 with PBS, filtered with 0.22 mm filters (Merck Millipore) and centrifuged in a *Optima™ TLX Preparative Ultracentrifuge, Beckman CoulterTM* at 150.000 g for 120 minutes at 4°C with a *TLA-55 Rotor* (*Beckman CoulterTM*) to pellet EVs. After supernatant was carefully removed, EV-containing pellets were stored at −80°C until use.

### Nanoparticle Tracking Analysis

Nanoparticle tracking analysis (NTA) was performed according to manufacturer’s instructions using a NanoSight NS300 system (Malvern Technologies, Malvern, UK) configured with 532 nm laser. All samples were diluted in filtered PBS to a final volume of 1 ml. Ideal measurement concentrations were found by pre-testing the ideal particle per frame value (20–100 particles/frame). Following settings were adjusted according to the manufacturer’s software manual. A syringe pump with constant flow injection was used and three videos of 60 s were captured and analyzed with NTA software version 3.2. From each video, the mean, mode, and median EVs size was used to calculate samples concentration expressed in nanoparticles/mL

### Protein quantification

We performed Bradford Assay to quantify protein concentration on our samples. The samples were added in a Bradford solution (BioRad Protein Assay 500-0006) 1:5 diluted in water and analysed by a spectrophotometer (Labsystem, Multiskan Ascent) at the wavelength of 595 nm. Furthermore, we analysed standard protein solutions to build a calibration line to discover the right protein concentration of our samples.

### SDS-PAGE and Western blot analysis

Treated EVs were added at Laemmli buffer and boiled for 5 minutes at 95□°C. Specifically, 10□μg of EVs were prepared in non-reducing conditions for tetraspanins detection, while 10□μg were used for soluble protein detection. Proteins were separated by SDS-PAGE (4-20%, Mini-Protean TGX Precast protein gel, Bio-Rad) and transferred onto a nitrocellulose membrane (BioRad, Trans-Blot Turbo). Nonspecific sites were saturated with a blocking solution for 1h (EveryBlot Blocking Buffer, BioRad). Membranes were incubated overnight at 4□°C with anti-CD9 (1:1000, BD Pharmingen), anti-CD63 (1:1000; BD Pharmingen,), anti-Alix (1:1000, Santa Cruz), anti-TSG101 (1:1000, Novus Bio) and anti-Apo1 (1:1000, Santa Cruz). After washing with T-TBS, membranes were incubated with the horseradish peroxidase-conjugated (Jackson ImmunoResearch) secondary antibodies diluted 1:3000 for 1 hour. After washing, the signal was detected using Bio-Rad Clarity Western ECL Substrate (Bio-Rad) and imaged using a Chemidoc XRS+ (BioRad).

### AFM sample preparation, imaging and morphometry

Borosilicate glass coverslips (Menzel Gläser GmbH, Germany) were first incubated for 1h in 2:1 H_2_SO_4_:H_2_O_2_(30% v/v) solution, then rinsed with ultrapure water, subjected to 3 x 30 minutes successive sonication cycles in acetone, isopropanol and ultrapure water, and finally dried under gentle nitrogen flow. Glass slides were then exposed for 5 minutes to air plasma and functionalized with (3-Aminopropyl)triethoxysilane (APTES) in vapor phase for 2h. Resuspended EVs solutions were diluted 1:100 with ultrapure water; 5 μl aliquots of the diluted solutions were then left to adsorb on APTES-functionalized slides for 30 minutes. AFM imaging was performed in PeakForce mode on a Multimode8 AFM microscope equipped with a type JV scanner and a sealed fluid cell (Bruker, USA). Image analysis was performed as described elsewhere^15^.

### EV array

Silicon slides (SVM, Sunnyvail, CA) were coated by MCP6 polymers (Lucidant Polymers) and spotted with BP membrane binding peptide (RPPGFSPRKG) synthesized as described in^19^. Printed slides were placed in a humid chamber overnight at room temperature. EVs samples wereincubated for 2 hours at particles concentration of 10^10^ particles/mL. Subsequently, the samples were removed and the slides were washed with washing buffer and incubated with anti-CD9-Biotin, anti-CD63-Biotin and anti-CD81-Biotin antibodies (Ancell) 0.1mg/mL for 1 hour. Then, the slide were incubated with Streptavidin-Cy3 (Jackson ImmunoResearch) 0.1mg/mL for 1 hour. Finally, slides were washed and dried and the analysis were performed by TECAN power scanner 50% laser intensity and 500% gain.

### miRNA isolation and retrotrascription

miRNAs were isolated from ultracentifuged EVs resuspended in 25 ul of phosphate buffered saline (PBS) using the Maxwell® RSC miRNA Plasma and Serum Kit (AS1680, Promega) following the manufacturer’s instruction. The RNA was eluted in 35μl of nuclease-free water.

cDNA was obtained using the TaqMan® MicroRNA Reverse Transcription kit (ThermoFisher) combined with TaqMan MicroRNA Assays (ThermoFisher). In particular, we used 5 ul of eluted RNA and 3 ul of primers specific for human miR-16 (assay ID 000391) and miR-21 (ID 000397). The reaction was performed with an initial incubation at 16°C for 30 min and a following step at 42°C for 30 min, finally, in order to terminate the RT step, a final incubation at 85°C for 5 min was succeeded.

### ddPCR reagents and cycling conditions

The miR-16-5p and miR-21-5p expression levels were performed by droplet digital PCR (ddPCR) using the QX100 ddPCR platform (Bio-Rad, Hercules, CA). The QX100 droplet generator was used to generate an emulsion of about 20,000 droplets. The volume of the PCR mix was 20 μL including 10 μL of ddPCR™ Supermix for Probes (No dUTP), 1 μL of probe (miR-16 or miR-21) and 5 ul of cDNA template. The droplet emulsion was thermally cycled on C1000 Touch Thermal Cycler (Bio-Rad) instrument. Cycling conditions were 95°C for 5 min, followed by 40 cycles of amplification (94°C for 30 s and 55°C for 1 min), ending with 98°C for 10 min, according to the manufacturer’s protocol. The concentration of the target was calculated automatically by the QuantaSoft™ software version 1.7.4 (Bio-Rad).

### EV-TRACK

We have submitted all relevant data of our experiments to the EV-TRACK knowledgebase (EV-TRACK ID: EV200180) (Van Deun J, et al. *EV-TRACK: transparent reporting and centralizing knowledge in extracellular vesicle research*. Nature methods. 2017;14(3):228-32).”

## Acknowledgements

Work partially funded from the European Union’s Horizon 2020 research and innovation programme under grant agreements No. 951768 (project MARVEL), No. 801367 (project EVfoundry), No. 952183 (project BOW), and Regione Lombardia&Fondazione Cariplo, grant n° 2018-1720 (project HYDROGEX).

## References

(1) Pastorino, B.; Touret, F.; Gilles, M.; de Lamballerie, X.; Charrel, R. N. Heat Inactivation of Different Types of SARS-CoV-2 Samples: What Protocols for Biosafety, Molecular Detection and Serological Diagnostics? Viruses 2020, 12 (7), 735. https://doi.org/10.3390/v12070735.

(2) Pastorino, B.; Touret, F.; Gilles, M.; Luciani, L.; de Lamballerie, X.; Charrel, R. N. Evaluation of Chemical Protocols for Inactivating SARS-CoV-2 Infectious Samples. Viruses 2020, 12 (6), 624. https://doi.org/10.3390/v12060624.

(3) Darnell, M. E. R.; Taylor, D. R. Evaluation of Inactivation Methods for Severe Acute Respiratory Syndrome Coronavirus in Noncellular Blood Products. Transfusion 2006. https://doi.org/10.1111/j.1537-2995.2006.00976.x.

(4) Hu, X.; Zhang, R.; An, T.; Li, Q.; Situ, B.; Ou, Z.; Wu, C.; Yang, B.; Tian, P.; Hu, Y.; Ping, B.; Wang, Q.; Zheng, L. Impact of Heat-Inactivation on the Detection of SARS-CoV-2 IgM and IgG Antibody by ELISA. Clinica Chimica Acta 2020, 509, 288–292. https://doi.org/10.1016/j.cca.2020.06.032.

(5) Hu, X.; An, T.; Situ, B.; Hu, Y.; Ou, Z.; Li, Q.; He, X.; Zhang, Y.; Tian, P.; Sun, D.; Rui, Y.; Wang, Q.; Ding, D.; Zheng, L. Heat Inactivation of Serum Interferes with the Immunoanalysis of Antibodies to SARS CoV 2. Journal of Clinical Laboratory Analysis 2020. https://doi.org/10.1002/jcla.23411.

(6) Nieuwland, R.; Falcón Pérez. J. M.; Théry, C.; Witwer, K. W. Rigor and Standardization of Extracellular Vesicle Research: Paving the Road towards Robustness. Journal of Extracellular Vesicles 2020, 10 (2). https://doi.org/10.1002/jev2.12037.

(7) Théry, C.; Witwer, K. W.; Aikawa, E.; Alcaraz, M. J.; Anderson, J. D.; Andriantsitohaina, R.; Antoniou, A.; Arab, T.; Archer, F.; Atkin-Smith, G. K.; Ayre, D. C.; Bach, J.-M. M.; Bachurski, D.; Baharvand, H.; Balaj, L.; Baldacchino, S.; Bauer, N. N.; Baxter, A. A.; Bebawy, M.; Beckham, C.; Bedina Zavec, A.; Benmoussa, A.; Berardi, A. C.; Bergese, P.; Bielska, E.; Blenkiron, C.; Bobis-Wozowicz, S.; Boilard, E.; Boireau, W.; Bongiovanni, A.; Borràs, F. E.; Bosch, S.; Boulanger, C. M.; Breakefield, X.; Breglio, A. M.; Brennan, M. Á.; Brigstock, D. R.; Brisson, A.; Broekman, M. L. D. L.; Bromberg, J. F.; Bryl-Górecka, P.; Buch, S.; Buck, A. H.; Burger, D.; Busatto, S.; Buschmann, D.; Bussolati, B.; Buzás, E. I.; Byrd, J. B.; Camussi, G.; Carter, D. R. F. R.; Caruso, S.; Chamley, L. W.; Chang, Y.-T. T.; Chaudhuri, A. D.; Chen, C.; Chen, S.; Cheng, L.; Chin, A. R.; Clayton, A.; Clerici, S. P.; Cocks, A.; Cocucci, E.; Coffey, R. J.; Cordeiro-da-Silva, A.; Couch, Y.; Coumans, F. A. A. W.; Coyle, B.; Crescitelli, R.; Criado, M. F.; D’Souza-Schorey, C.; Das, S.; de Candia, P.; de Santana, E. F.; de Wever, O.; del Portillo, H. A.; Demaret, T.; Deville, S.; Devitt, A.; Dhondt, B.; di Vizio, D.; Dieterich, L. C.; Dolo, V.; Dominguez Rubio, A. P.; Dominici, M.; Dourado, M. R.; Driedonks, T. A. A. P.; Duarte, F.v.; Duncan, H. M.; Eichenberger, R. M.; Ekström, K.; el Andaloussi, S.; Elie-Caille, C.; Erdbrügger, U.; Falcón-Pérez, J. M.; Fatima, F.; Fish, J. E.; Flores-Bellver, M.; Försönits, A.; Frelet-Barrand, A.; Fricke, F.; Fuhrmann, G.; Gabrielsson, S.; Gámez-Valero, A.; Gardiner, C.; Gärtner, K.; Gaudin, R.; Gho, Y. S.; Giebel, B.; Gilbert, C.; Gimona, M.; Giusti, I.; Goberdhan, D. C. I. C.; Görgens, A.; Gorski, S. M.; Greening, D. W.; Gross, J. C.; Gualerzi, A.; Gupta, G. N.; Gustafson, D.; Handberg, A.; Haraszti, R. A.; Harrison, P.; Hegyesi, H.; Hendrix, A.; Hill, A. F.; Hochberg, F. H.; Hoffmann, K. F.; Holder, B.; Holthofer, H.; Hosseinkhani, B.; Hu, G.; Huang, Y.; Huber, V.; Hunt, S.; Ibrahim, A. G.-E. E.; Ikezu, T.; Inal, J. M.; Isin, M.; Ivanova, A.; Jackson, H. K.; Jacobsen, S.; Jay, S. M.; Jayachandran, M.; Jenster, G.; Jiang, L.; Johnson, S. M.; Jones, J. C.; Jong, A.; Jovanovic-Talisman, T.; Jung, S.; Kalluri, R.; Kano, S. ichi; Kaur, S.; Kawamura, Y.; Keller, E. T.; Khamari, D.; Khomyakova, E.; Khvorova, A.; Kierulf, P.; Kim, K. P.; Kislinger, T.; Klingeborn, M.; Klinke, D. J.; Kornek, M.; Kosanović, M. M.; Kovács, Á. F.; Krämer-Albers, E.-M. M.; Krasemann, S.; Krause, M.; Kurochkin, I.v.; Kusuma, G. D.; Kuypers, S.; Laitinen, S.; Langevin, S. M.; Languino, L. R.; Lannigan, J.; Lässer, C.; Laurent, L. C.; Lavieu, G.; Lázaro-Ibáñez, E.; le Lay, S.; Lee, M.-S. S.; Lee, Y. X. F.; Lemos, D. S.; Lenassi, M.; Leszczynska, A.; Li, I. T. T. S.; Liao, K.; Libregts, S. F.; Ligeti, E.; Lim, R.; Lim, S. K.; Linē, A.; Linnemannstöns, K.; Llorente, A.; Lombard, C. A.; Lorenowicz, M. J.; Lörincz, Á. M.; Lötvall, J.; Lovett, J.; Lowry, M. C.; Loyer, X.; Lu, Q.; Lukomska, B.; Lunavat, T. R.; Maas, S. L. L. N.; Malhi, H.; Marcilla, A.; Mariani, J.; Mariscal, J.; Martens-Uzunova, E. S.; Martin-Jaular, L.; Martinez, M. C.; Martins, V. R.; Mathieu, M.; Mathivanan, S.; Maugeri, M.; McGinnis, L. K.; McVey, M. J.; Meckes, D. G.; Meehan, K. L.; Mertens, I.; Minciacchi, V. R.; Möller, A.; Møller Jørgensen, M.; Morales-Kastresana, A.; Morhayim, J.; Mullier, F.; Muraca, M.; Musante, L.; Mussack, V.; Muth, D. C.; Myburgh, K. H.; Najrana, T.; Nawaz, M.; Nazarenko, I.; Nejsum, P.; Neri, C.; Neri, T.; Nieuwland, R.; Nimrichter, L.; Nolan, J. P.; Nolte-’t Hoen, E. N. M. N.; Noren Hooten, N.; O’Driscoll, L.; O’Grady, T.; O’Loghlen, A.; Ochiya, T.; Olivier, M.; Ortiz, A.; Ortiz, L. A.; Osteikoetxea, X.; Ostegaard, O.; Ostrowski, M.; Park, J.; Pegtel, D. M.; Peinado, H.; Perut, F.; Pfaffl, M. W.; Phinney, D. G.; Pieters, B. C. C. H.; Pink, R. C.; Pisetsky, D. S.; Pogge von Strandmann, E.; Polakovicova, I.; Poon, I. K. K. H.; Powell, B. H.; Prada, I.; Pulliam, L.; Quesenberry, P.; Radeghieri, A.; Raffai, R. L.; Raimondo, S.; Rak, J.; Ramirez, M. I.; Raposo, G.; Rayyan, M. S.; Regev-Rudzki, N.; Ricklefs, F. L.; Robbins, P. D.; Roberts, D. D.; Rodrigues, S. C.; Rohde, E.; Rome, S.; Rouschop, K. M. M. A.; Rughetti, A.; Russell, A. E.; Saá, P.; Sahoo, S.; Salas-Huenuleo, E.; Sánchez, C.; Saugstad, J. A.; Saul, M. J.; Schiffelers, R. M.; Schneider, R.; Schøyen, T. H.; Scott, A.; Shahaj, E.; Sharma, S.; Shatnyeva, O.; Shekari, F.; Shelke, G. V.; Shetty, A. K.; Shiba, K.; Siljander, P. R. M. R.-M.; Silva, A. M.; Skowronek, A.; Snyder, O. L.; Soares, R. P.; Sódar, B. W.; Soekmadji, C.; Sotillo, J.; Stahl, P. D.; Stoorvogel, W.; Stott, S. L.; Strasser, E. F.; Swift, S.; Tahara, H.; Tewari, M.; Timms, K.; Tiwari, S.; Tixeira, R.; Tkach, M.; Toh, W. S.; Tomasini, R.; Torrecilhas, A. C.; Tosar, J. P.; Toxavidis, V.; Urbanelli, L.; Vader, P.; van Balkom, B. W. M. W.; van der Grein, S. G.; van Deun, J.; van Herwijnen, M. J. C. J.; van Keuren-Jensen, K.; van Niel, G.; van Royen, M. E.; van Wijnen, A. J.; Vasconcelos, M. H.; Vechetti, I. J.; Veit, T. D.; Vella, L. J.; Velot, É.; Verweij, F. J.; Vestad, B.; Viñas, J. L.; Visnovitz, T.; Vukman, K.v.; Wahlgren, J.; Watson, D. C.; Wauben, M. H. H. M.; Weaver, A.; Webber, J. P.; Weber, V.; Wehman, A. M.; Weiss, D. J.; Welsh, J. A.; Wendt, S.; Wheelock, A. M.; Wiener, Z.; Witte, L.; Wolfram, J.; Xagorari, A.; Xander, P.; Xu, J.; Yan, X.; Yáñez-Mó, M.; Yin, H.; Yuana, Y.; Zappulli, V.; Zarubova, J.; Žėkas, V.; Zhang, J. ye; Zhao, Z.; Zheng, L.; Zheutlin, A. R.; Zickler, A. M.; Zimmermann, P.; Zivkovic, A. M.; Zocco, D.; Zuba-Surma, E. K.; Datta Chaudhuri, A.; de Candia, P.; de Santana, E. F.; de Wever, O.; del Portillo, H. A.; Demaret, T.; Deville, S.; Devitt, A.; Dhondt, B.; di Vizio, D.; Dieterich, L. C.; Dolo, V.; Dominguez Rubio, A. P.; Dominici, M.; Dourado, M. R.; Driedonks, T. A. A. P.; Duarte, F.v.; Duncan, H. M.; Eichenberger, R. M.; Ekström, K.; el Andaloussi, S.; Elie-Caille, C.; Erdbrügger, U.; Falcón-Pérez, J. M.; Fatima, F.; Fish, J. E.; Flores-Bellver, M.; Försönits, A.; Frelet-Barrand, A.; Fricke, F.; Fuhrmann, G.; Gabrielsson, S.; Gámez-Valero, A.; Gardiner, C.; Gärtner, K.; Gaudin, R.; Gho, Y. S.; Giebel, B.; Gilbert, C.; Gimona, M.; Giusti, I.; Goberdhan, D. C. I. C.; Görgens, A.; Gorski, S. M.; Greening, D. W.; Gross, J. C.; Gualerzi, A.; Gupta, G. N.; Gustafson, D.; Handberg, A.; Haraszti, R. A.; Harrison, P.; Hegyesi, H.; Hendrix, A.; Hill, A. F.; Hochberg, F. H.; Hoffmann, K. F.; Holder, B.; Holthofer, H.; Hosseinkhani, B.; Hu, G.; Huang, Y.; Huber, V.; Hunt, S.; Ibrahim, A. G.E. E.; Ikezu, T.; Inal, J. M.; Isin, M.; Ivanova, A.; Jackson, H. K.; Jacobsen, S.; Jay, S. M.; Jayachandran, M.; Jenster, G.; Jiang, L.; Johnson, S. M.; Jones, J. C.; Jong, A.; Jovanovic-Talisman, T.; Jung, S.; Kalluri, R.; Kano, S. ichi; Kaur, S.; Kawamura, Y.; Keller, E. T.; Khamari, D.; Khomyakova, E.; Khvorova, A.; Kierulf, P.; Kim, K. P.; Kislinger, T.; Klingeborn, M.; Klinke, D. J.; Kornek, M.; Kosanović, M. M.; Kovács, Á. F.; Krämer-Albers, E.-M. M.; Krasemann, S.; Krause, M.; Kurochkin, I. v.; Kusuma, G. D.; Kuypers, S.; Laitinen, S.; Langevin, S. M.; Languino, L. R.; Lannigan, J.; Lässer, C.; Laurent, L. C.; Lavieu, G.; Lázaro-Ibáñez, E.; le Lay, S.; Lee, M.-S. S.; Lee, Y. X. F.; Lemos, D. S.; Lenassi, M.; Leszczynska, A.; Li, I. T. T. S.; Liao, K.; Libregts, S. F.; Ligeti, E.; Lim, R.; Lim, S. K.; Linē, A.; Linnemannstöns, K.; Llorente, A.; Lombard, C. A.; Lorenowicz, M. J.; Lörincz, Á. M.; Lötvall, J.; Lovett, J.; Lowry, M. C.; Loyer, X.; Lu, Q.; Lukomska, B.; Lunavat, T. R.; Maas, S. L. L. N.; Malhi, H.; Marcilla, A.; Mariani, J.; Mariscal, J.; Martens-Uzunova, E. S.; Martin-Jaular, L.; Martinez, M. C.; Martins, V. R.; Mathieu, M.; Mathivanan, S.; Maugeri, M.; McGinnis, L. K.; McVey, M. J.; Meckes, D. G.; Meehan, K. L.; Mertens, I.; Minciacchi, V. R.; Möller, A.; Møller Jørgensen, M.; Morales-Kastresana, A.; Morhayim, J.; Mullier, F.; Muraca, M.; Musante, L.; Mussack, V.; Muth, D. C.; Myburgh, K. H.; Najrana, T.; Nawaz, M.; Nazarenko, I.; Nejsum, P.; Neri, C.; Neri, T.; Nieuwland, R.; Nimrichter, L.; Nolan, J. P.; Nolte-’t Hoen, E. N. M. N.; Noren Hooten, N.; O’Driscoll, L.; O’Grady, T.; O’Loghlen, A.; Ochiya, T.; Olivier, M.; Ortiz, A.; Ortiz, L. A.; Osteikoetxea, X.; Østergaard, O.; Ostrowski, M.; Park, J.; Pegtel, D. M.; Peinado, H.; Perut, F.; Pfaffl, M. W.; Phinney, D. G.; Pieters, B. C. C. H.; Pink, R. C.; Pisetsky, D. S.; Pogge von Strandmann, E.; Polakovicova, I.; Poon, I. K. K. H.; Powell, B. H.; Prada, I.; Pulliam, L.; Quesenberry, P.; Radeghieri, A.; Raffai, R. L.; Raimondo, S.; Rak, J.; Ramirez, M. I.; Raposo, G.; Rayyan, M. S.; Regev-Rudzki, N.; Ricklefs, F. L.; Robbins, P. D.; Roberts, D. D.; Rodrigues, S. C.; Rohde, E.; Rome, S.; Rouschop, K. M. M. A.; Rughetti, A.; Russell, A. E.; Saá, P.; Sahoo, S.; Salas-Huenuleo, E.; Sánchez, C.; Saugstad, J. A.; Saul, M. J.; Schiffelers, R. M.; Schneider, R.; Schøyen, T. H.; Scott, A.; Shahaj, E.; Sharma, S.; Shatnyeva, O.; Shekari, F.; Shelke, G. V.; Shetty, A. K.; Shiba, K.; Siljander, P. R. M. R.-M.; Silva, A. M.; Skowronek, A.; Snyder, O. L.; Soares, R. P.; Sódar, B. W.; Soekmadji, C.; Sotillo, J.; Stahl, P. D.; Stoorvogel, W.; Stott, S. L.; Strasser, E. F.; Swift, S.; Tahara, H.; Tewari, M.; Timms, K.; Tiwari, S.; Tixeira, R.; Tkach, M.; Toh, W. S.; Tomasini, R.; Torrecilhas, A. C.; Tosar, J. P.; Toxavidis, V.; Urbanelli, L.; Vader, P.; van Balkom, B. W. M. W.; van der Grein, S. G.; van Deun, J.; van Herwijnen, M. J. C. J.; van Keuren-Jensen, K.; van Niel, G.; van Royen, M. E.; van Wijnen, A. J.; Vasconcelos, M. H.; Vechetti, I. J.; Veit, T. D.; Vella, L. J.; Velot, É.; Verweij, F. J.; Vestad, B.; Viñas, J. L.; Visnovitz, T.; Vukman, K. v.; Wahlgren, J.; Watson, D. C.; Wauben, M. H. H. M.; Weaver, A.; Webber, J. P.; Weber, V.; Wehman, A. M.; Weiss, D. J.; Welsh, J. A.; Wendt, S.; Wheelock, A. M.; Wiener, Z.; Witte, L.; Wolfram, J.; Xagorari, A.; Xander, P.; Xu, J.; Yan, X.; Yáñez-Mó, M.; Yin, H.; Yuana, Y.; Zappulli, V.; Zarubova, J.; Žėkas, V.; Zhang, J. ye; Zhao, Z.; Zheng, L.; Zheutlin, A. R.; Zickler, A. M.; Zimmermann, P.; Zivkovic, A. M.; Zocco, D.; Zuba-Surma, E. K. Minimal Information for Studies of Extracellular Vesicles 2018 (MISEV2018): A Position Statement of the International Society for Extracellular Vesicles and Update of the MISEV2014 Guidelines. Journal of Extracellular Vesicles 2018, 7 (1), 1535750. https://doi.org/10.1080/20013078.2018.1535750.

(8) Schulz, E.; Karagianni, A.; Koch, M.; Fuhrmann, G. Hot EVs – How Temperature Affects Extracellular Vesicles. European Journal of Pharmaceutics and Biopharmaceutics 2020, 146, 55–63. https://doi.org/10.1016/j.ejpb.2019.11.010.

(9) Osteikoetxea, X.; Sódar, B.; Németh, A.; Szabó-Taylor, K.; Pálóczi, K.; Vukman, K.v.; Tamási, V.; Balogh, A.; Kittel, Á.; Pállinger, É.; Buzás, E. I. Differential Detergent Sensitivity of Extracellular Vesicle Subpopulations. Organic and Biomolecular Chemistry 2015. https://doi.org/10.1039/c5ob01451d.

(10) Hong, C.-S.; Funk, S.; Muller, L.; Boyiadzis, M.; Whiteside, T. L. Isolation of Biologically Active and Morphologically Intact Exosomes from Plasma of Patients with Cancer. Journal of Extracellular Vesicles 2016, 5 (1), 29289. https://doi.org/10.3402/jev.v5.29289.

(11) Lobb, R. J.; Becker, M.; Wen Wen, S.; Wong, C. S. F.; Wiegmans, A. P.; Leimgruber, A.; Möller, A. Optimized Exosome Isolation Protocol for Cell Culture Supernatant and Human Plasma. Journal of Extracellular Vesicles 2015, 4 (1), 27031. https://doi.org/10.3402/jev.v4.27031.

(12) Zhang, X.; Borg, E. G. F.; Liaci, A. M.; Vos, H. R.; Stoorvogel, W. A Novel Three Step Protocol to Isolate Extracellular Vesicles from Plasma or Cell Culture Medium with Both High Yield and Purity. Journal of Extracellular Vesicles 2020, 9 (1), 1791450. https://doi.org/10.1080/20013078.2020.1791450.

(13) Karimi, N.; Cvjetkovic, A.; Jang, S. C.; Crescitelli, R.; Hosseinpour Feizi, M. A.; Nieuwland, R.; Lötvall, J.; Lässer, C. Detailed Analysis of the Plasma Extracellular Vesicle Proteome after Separation from Lipoproteins. Cellular and Molecular Life Sciences 2018. https://doi.org/10.1007/s00018-018-2773-4.

(14) Osteikoetxea, X.; Sódar, B.; Németh, A.; Szabó-Taylor, K.; Pálóczi, K.; Vukman, K.v.; Tamási, V.; Balogh, A.; Kittel, Á.; Pállinger, É.; Buzás, E. I. Differential Detergent Sensitivity of Extracellular Vesicle Subpopulations. Organic & Biomolecular Chemistry 2015, 13 (38), 9775–9782. https://doi.org/10.1039/C5OB01451D.

(15) Ridolfi, A.; Brucale, M.; Montis, C.; Caselli, L.; Paolini, L.; Borup, A.; Boysen, A. T.; Loria, F.; van Herwijnen, M. J. C.; Kleinjan, M.; Nejsum, P.; Zarovni, N.; Wauben, M. H. M.; Berti, D.; Bergese, P.; Valle, F. AFM-Based High-Throughput Nanomechanical Screening of Single Extracellular Vesicles. Analytical Chemistry 2020, 92 (15), 10274–10282. https://doi.org/10.1021/acs.analchem.9b05716.

(16) Ridolfi, A.; Caselli, L.; Montis, C.; Mangiapia, G.; Berti, D.; Brucale, M.; Valle, F. Gold Nanoparticles Interacting with Synthetic Lipid Rafts: An AFM Investigation. Journal of Microscopy 2020, 280 (3), 194–203. https://doi.org/10.1111/jmi.12910.

(17) Rikkert, L. G.; Nieuwland, R.; Terstappen, L. W. M. M.; Coumans, F. A. W. Quality of Extracellular Vesicle Images by Transmission Electron Microscopy Is Operator and Protocol Dependent. Journal of Extracellular Vesicles 2019, 8 (1), 1555419. https://doi.org/10.1080/20013078.2018.1555419.

(18) Jørgensen, M.; Bæk, R.; Pedersen, S.; Søndergaard, E. K. L. L.; Kristensen, S. R.; Varming, K. Extracellular Vesicle (EV) Array: Microarray Capturing of Exosomes and Other Extracellular Vesicles for Multiplexed Phenotyping. Journal of Extracellular Vesicles 2013, 2 (1), 20920. https://doi.org/10.3402/jev.v2i0.20920.

(19) Gori, A.; Romanato, A.; Greta, B.; Strada, A.; Gagni, P.; Frigerio, R.; Brambilla, D.; Vago, R.; Galbiati, S.; Picciolini, S.; Bedoni, M.; Daaboul, G. G.; Chiari, M.; Cretich, M. Membrane-Binding Peptides for Extracellular Vesicles on-Chip Analysis. Journal of Extracellular Vesicles 2020. https://doi.org/10.1080/20013078.2020.1751428.

(20) Cretich, M.; di Carlo, G.; Longhi, R.; Gotti, C.; Spinella, N.; Coffa, S.; Galati, C.; Renna, L.; Chiari, M. High Sensitivity Protein Assays on Microarray Silicon Slides. Analytical Chemistry 2009, 81 (13), 5197–5203. https://doi.org/10.1021/ac900658c.

(21) Cretich, M.; Bagnati, M.; Damin, F.; Sola, L.; Chiari, M. Overcoming Mass Transport Limitations to Achieve Femtomolar Detection Limits on Silicon Protein Microarrays. Analytical Biochemistry 2011, 418 (1), 164–166. https://doi.org/10.1016/j.ab.2011.07.004.

(22) Michell, D. L.; Vickers, K. C. Lipoprotein Carriers of MicroRNAs. Biochimica et Biophysica Acta (BBA) -Molecular and Cell Biology of Lipids 2016, 1861 (12), 2069–2074. https://doi.org/10.1016/j.bbalip.2016.01.011.

(23) Karlsen, T. A.; Aae, T. F.; Brinchmann, J. E. Robust Profiling of MicroRNAs and IsomiRs in Human Plasma Exosomes across 46 Individuals. Scientific Reports 2019, 9 (1), 19999. https://doi.org/10.1038/s41598-019-56593-7.

(24) Axmann, M.; Meier, S.; Karner, A.; Strobl, W.; Stangl, H.; Plochberger, B. Serum and Lipoprotein Particle MiRNA Profile in Uremia Patients. Genes 2018, 9 (11), 533. https://doi.org/10.3390/genes9110533.

(25) Weber, J. A.; Baxter, D. H.; Zhang, S.; Huang, D. Y.; Huang, K. H.; Lee, M. J.; Galas, D. J.; Wang, K. The MicroRNA Spectrum in 12 Body Fluids. Clinical Chemistry 2010. https://doi.org/10.1373/clinchem.2010.147405.

(26) Borzi, C.; Calzolari, L.; Ferretti, A. M.; Caleca, L.; Pastorino, U.; Sozzi, G.; Fortunato, O. C-Myc Shuttled by Tumour-Derived Extracellular Vesicles Promotes Lung Bronchial Cell Proliferation through MiR-19b and MiR-92a. Cell Death & Disease 2019, 10 (10), 759. https://doi.org/10.1038/s41419-019-2003-5.

